# Cholesterol-Dependent Membrane Deformation by Metastable Viral Capsids Facilitates Entry

**DOI:** 10.1101/2024.01.10.575085

**Authors:** Mengchi Jiao, Pranav Danthi, Yan Yu

## Abstract

Non-enveloped viruses employ unique entry mechanisms to breach and infect host cells. Understanding these mechanisms is crucial for developing antiviral strategies. Prevailing perspective suggests that non-enveloped viruses release membrane lytic peptides to breach host membranes. However, the precise involvement of the viral capsid in this entry remains elusive. Our study presents direct observations elucidating the dynamically distinctive steps through which metastable reovirus capsids disrupt host lipid membranes as they uncoat into partially hydrophobic intermediate particles. Using both live cells and model membrane systems, our key finding is that reovirus capsids actively deform and permeabilize lipid membranes in a cholesterol-dependent process. Unlike membrane lytic peptides, these metastable viral capsids induce more extensive membrane perturbations, including budding, bridging between adjacent membranes, and complete rupture. Notably, cholesterol enhances subviral particle adsorption, resulting in the formation of pores equivalent to the capsid size. This cholesterol dependence is attributed to the lipid condensing effect, particularly prominent at intermediate cholesterol level. Furthermore, our results reveal a positive correlation between membrane disruption extent and efficiency of viral variants in establishing infection. This study unveils the crucial role of capsid-lipid interaction in non-enveloped virus entry, providing new insights into how cholesterol homeostasis influences virus infection dynamics.

## Introduction

The entry of viruses into host cells is a crucial gateway for infection, influencing subsequent replication and pathogenesis. Understanding the interactions involved in virus-host cell membrane encounter not only sheds light on the fundamental biology of viruses but also informs the development of targeted antiviral therapies for effective control and mitigation of infectious diseases. Most viruses, both non-enveloped and enveloped viruses, initiate infection by attaching to host cells via receptors and conclude by delivering genomic material into the host cytoplasm for replication. In the case of enveloped viruses, fusion between the viral and host membranes is the typical mechanism for delivering the viral genomic payload ^1^. Molecular details of the fusion events for a variety of enveloped viruses are conserved and are well understood ^2-5^. In contrast, somewhat surprisingly, the membrane penetration mechanisms of non-enveloped viruses remain less understood.

Ensemble-average biochemistry assays have revealed a conserved general strategy for membrane penetration among non-enveloped viruses spanning diverse families, such as adenoviruses, picornaviruses, and reoviruses ^6^. Lacking a lipid bilayer shell, the naked capsids of these viruses undergo conformational changes upon encountering stimuli such as pH and proteases in the correct host environment. This leads to the release of membrane-active amphiphilic or hydrophobic peptides. The peptides alter the integrity of host membranes through mechanisms that remain unresolved, allowing the delivery of the viral genomic material. Studies have shown that peptides from adenovirus rupture the host membrane ^7^, whereas those from picornaviruses and reoviruses induce pores instead ^8-12^. However, as straightforward as this general mechanism of membrane perforation appears, it cannot explain how some non-enveloped viruses, such as reovirus and bluetongue virus, can transport their capsid core of approximately 70 nm across pores that were measured to be merely a few nanometers ^13, 14^. Recent studies suggest that the released peptides not only form pores in the membrane but also cooperate with host lipids to recruit virus capsids ^15, 16^. However, the process by which this and other non-enveloped viruses utilize their interaction with lipids and the released peptides to coordinate membrane disruption for productive cell entry remains poorly understood.

In this study, we investigate the dynamic interaction of reovirus, a model non-enveloped virus, with lipid membranes and, importantly, how this interaction leads to membrane deformation. Reoviruses undergo stepwise disassembly to complete cell entry. As shown in **Figure 1A**, virions are initially digested by proteases, which can occur either extracellularly or within endosomes, generating a metastable entry intermediate referred to as the infectious subvirion particle (ISVP)^17^. During this initial conversion step, virions lose the σ3 protein on the capsid, and the μ1 protein is cleaved into two particle-associated fragments, μ1δ and ϕ ^18^. The ISVPs then undergo further conformational changes to generate ISVP* ^19^. During this conversion step, μ1δ on the ISVP surface is autocleaved to expose the amphiphilic μ1 peptide, μ1N (**Figure. 1B**). As the ISVPs uncoat to become ISVP*s, μ1N is released along with ϕ ^20, 21^. Our previous studies, using ensemble-average biochemical assays, have shown that this ISVP-to-ISVP* conversion is facilitated by lipids of the host cell membrane ^15, 16^. However, interactions between the dynamic, metastable viral capsids and lipids have not been observed directly. Such real-time visualization is expected to provide crucial insights into understanding how the viral capsid-lipid interaction leads to membrane disruption for viral entry.

**Figure 1.**
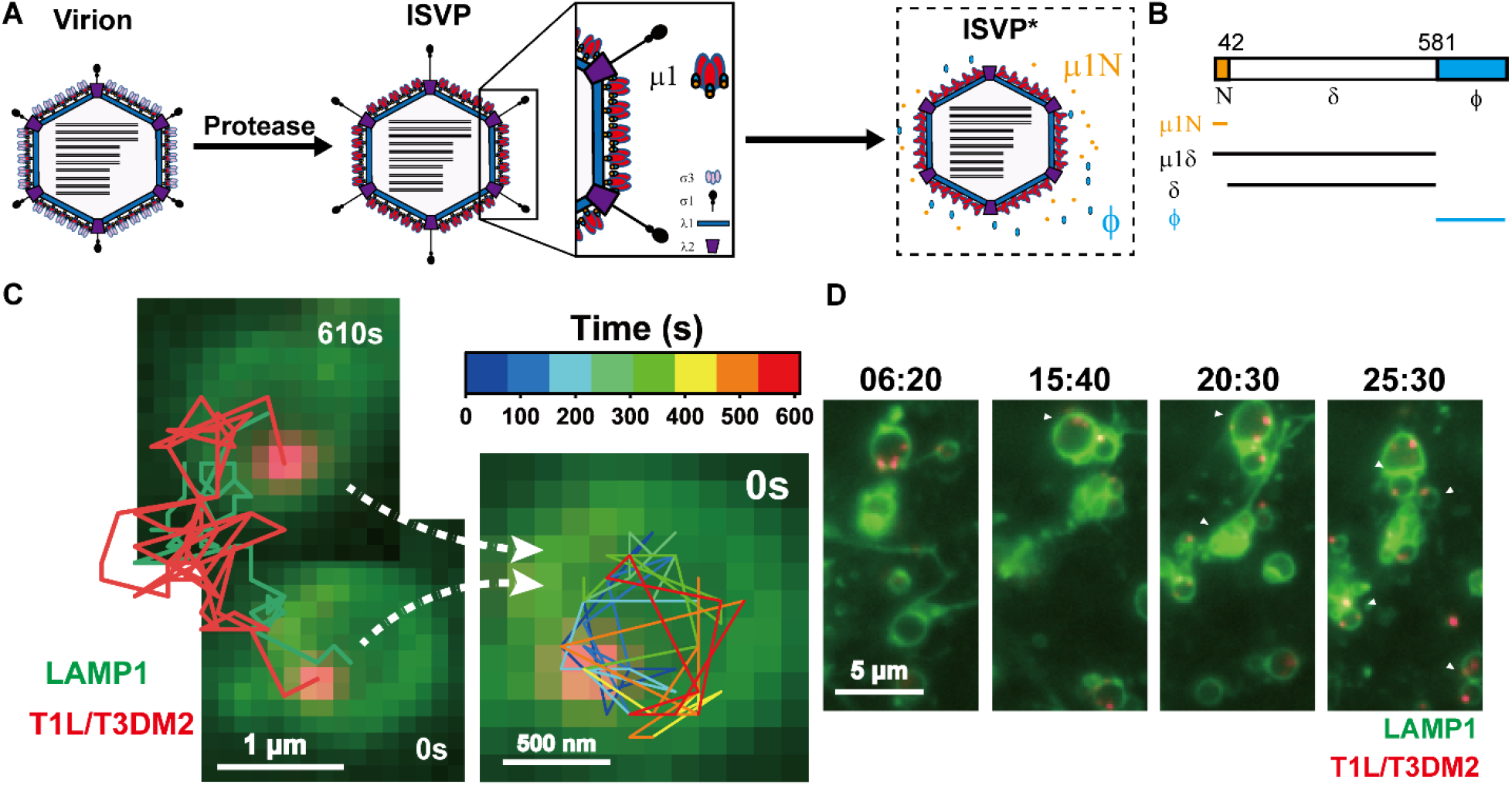
T1L/T3DM2 reovirus induces deformation of endosomal membrane in Vero 76 cells. (A) Schematic illustration of the *in vitro* structural transformation of reovirus virion to intermediate Infectious Subvirion particle (ISVP) and finally ISVP*. (B) Schematic illustration of cleavage fragments of the μ1 protein formed during the disassembly of ISVP capsid. (C) TIRF microscopy time-lapse fluorescence images and line plots showing the trajectory of a T1L/T3DM2 reovirus (shown in red) within endolysosomes marked by LAMP1-GFP (shown in green). The trajectories show the movement of reovirus along the inner membrane of the endolysosome. (D) TIRF microscopy time-lapse fluorescence images showing the bridging of LAMP1-positive endolysosomes encapsulating T1L/T3DM2 reoviruses.

In this study, we directly observe the process during which reovirus capsids interact with and disrupt host membranes, both in live cells and model membrane systems. A key finding from our research is that metastable reovirus capsids, as they disassemble to become partially hydrophobic particles, deform and permeabilize lipid membranes through a cholesterol-dependent mechanism. The viral capsids, not the released peptides, induce membrane budding and bridging between adjacent membranes, in addition to some membrane rupture. The presence of cholesterol promotes the adsorption of ISVPs on lipid membranes and the formation of pores that are as large as the viral capsid. This cholesterol dependence is attributed to the lipid condensing effect, and it is most prominent at an intermediate level of cholesterol. The extent of membrane disruption is positively correlated with the infection efficiency of viral variants; viral capsids that induce more pronounced membrane deformation and rupture are able to infect cells more readily. Our results unveil the process by which the metastable capsid of non-enveloped viruses interacts with lipid membranes and highlight the distinct role of this interaction in facilitating viral cell entry.

## Results and Discussion

### Live-cell imaging of endosomal deformation induced by reoviruses

We first investigated the interaction between reovirus capsids and endosomal membranes in live cells. Virions are known to be internalized into endosomes during entry after receptor engagement. Endosomal proteases disassemble particles to form ISVPs. Based on the evidence that ISVP to ISVP conversion is needed to cross the membrane, we expect that ISVPs and ISVP*s exist within endolysosome compartments ^22^. In the experiments, we transiently transfected Vero 76 cells with a lysosomal-associated membrane protein 1 (LAMP1) GFP construct and then infected cells with reovirus (T1L/T3DM2). LAMP1-GFP serves as a marker for endolysosomes and lysosomes. The viral capsids were fluorescently labeled on the surface for imaging. Fluorescence images revealed that the viral capsids were encapsulated inside membrane compartments marked by LAMP1-GFP (Figure. 1 C, D). Intriguingly, these viral capsids moved along the inner membrane wall of many large-sized endosomes (Figure. 1B; Video S1). Concurrently, the virus-encapsulating endolysosomes gradually became bridged to one another (Figure. 1 D; Video S2). This membrane bridging bears morphological similarities to the aggregation of vesicles induced by membrane-active proteins and peptides ^23-25^. Given that both the metastable viral capsid of reoviruses and the released μ1 peptides interact with lipid membranes, we postulated that either of capsid or the peptide induces membrane instability, leading to subsequent membrane bridging. As live cell endosomes are too complex to test this hypothesis, we opted for experiments using reconstituted model membranes. An additional advantage of the model membrane approach is that we were able to directly investigate capsid-lipid membrane interaction without the interfering effect from receptor binding.

### Vesicle membrane deformations induced by viral capsids during uncoating

We used giant unilamellar vesicles (GUVs) as our model membrane system (Figure. 2 A). Their sizes range from a few to tens of micrometers, a good representation of the large endosomes we observed in cells. The large size of GUVs also facilitates the direct imaging of dynamic interactions between reoviruses and lipids through fluorescence microscopy. In these experiments, GUV membranes were composed of 67 mol% 1,2-dioleoyl-sn-glycero-3-phosphocholine (DOPC), 33 mol% 1,2-dioleoyl-sn-glycero-3-phosphoethanolamine (DOPE), and a trace amount (0.2 mol%) of fluorescently labeled lipid, BODIPY-PE. We chose this lipid composition because the 2:1 molar ratio of PC and PE is representative of the lipid compositions of early or late endosomal membranes, and was shown to promote ISVP to ISVP* conversion in our previous study ^15^.

**Figure 2.**
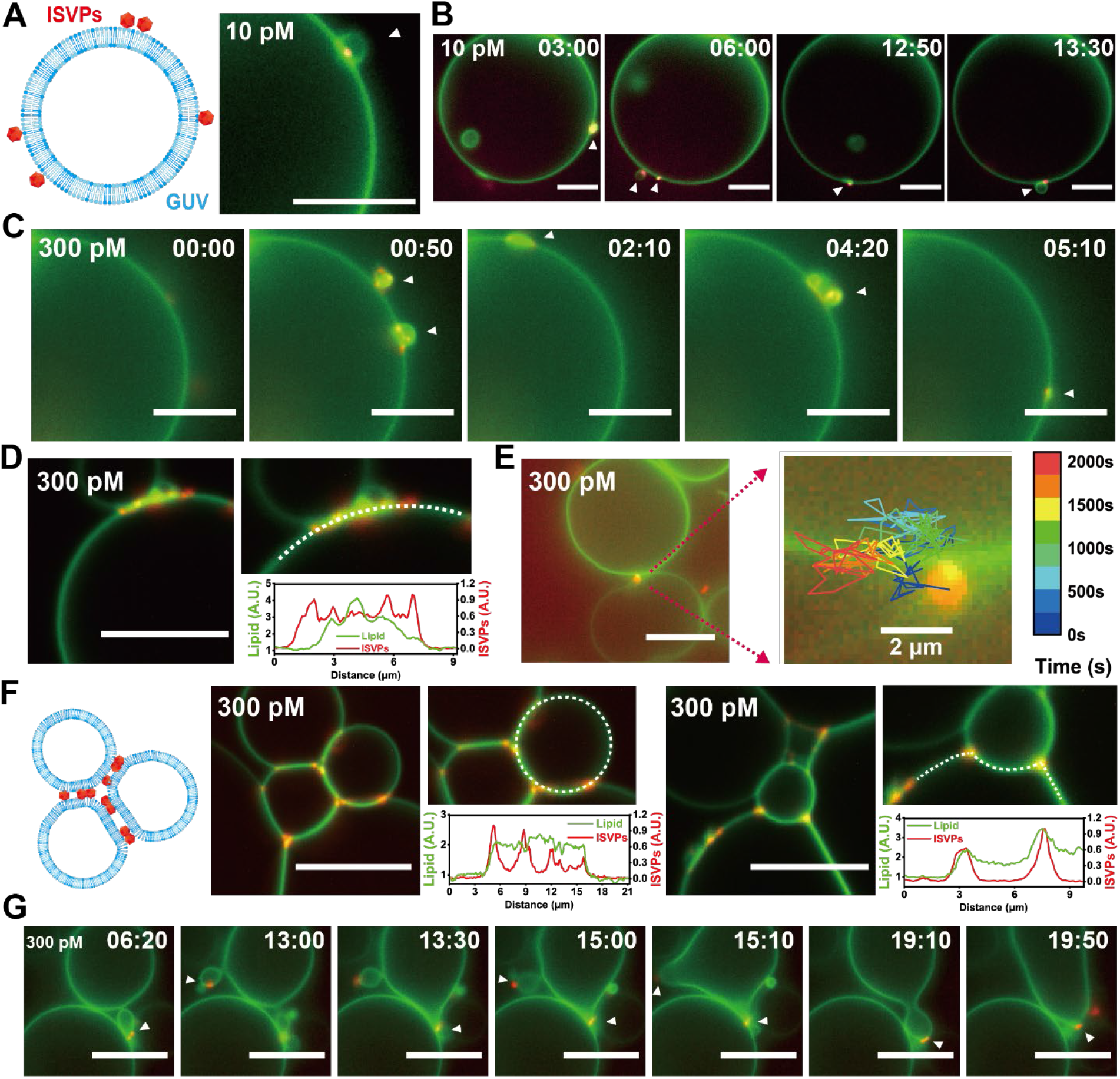
T1L ISVPs induced membrane deformation and bridging between GUVs. (A) Schematic illustration of reovirus ISVPs interaction with giant unilamellar vesicles (GUVs) and epifluorescence image showing the adsorption of T1L ISVPs (shown in red) and the membrane budding on a GUV (shown in green). T1L ISVP concentration was 10 pM. (B, C) Epifluorescence time-lapse images showing the diffusion of T1L ISVPs along the GUV surface and their colocalization with membrane budding (indicated by white arrows). ISVP concentration 10 pM in (B) and 300 pM in (C). (D) Rescan confocal microscopy images and line-scan plots showing the colocalization between T1L ISVPs (300 pM) and the membrane budding. (E) Epifluorescence images showing the bridging of two adjacent GUVs by T1L ISVPs (300 pM). The trajectory, color-coded with time, shows the confinement of ISVPs at the junction between two GUVs. (F) Schematic illustration and Rescan confocal microscopy images showing the extensive membrane bridging induced by T1L ISVPs (300 pM). Line-scan intensity plots show the localization of T1L ISVPs and the formation of two membranes after the bridging of GUVs. (G) Epifluorescence time-lapse images showing the fusion of GUVs induced by 300 pM T1L ISVPs. Scale bars in all images, 10 μm.

Because virions are not expected to encounter proteases and disassemble to form ISVPs in this minimalist system, we generated ISVPs from virions *in vitro* before incubating them with GUVs. We first added 10 pM of CF568 labeled type 1 Lang (T1L) ISVPs to the GUVs and imaged their dynamic interactions. The samples were kept at 37 °C during imaging to promote the ISVP to ISVP* conversion and the capsid-lipid interactions. Time-lapse fluorescence images revealed that some T1L ISVPs rapidly adsorbed on the membrane of GUVs (Figure. 2 A). After this, a small fraction (≈ 10%) of the GUVs ruptured to form lipid bilayers on the glass bottom of imaging chambers, while most GUVs remained in the solution but exhibited membrane deformations. For the GUVs that remained in the solution, we observed ISVPs “surfing” along the GUV membrane and gradually aggregating together, characterized by much higher fluorescence intensity than that of single ISVPs. After a few minutes of interaction, most ISVPs were seen to colocalize with membrane budding – protrusion of small vesicles (Figure. 2B, Video S3). This membrane deformation was dependent on the ISVP concentration. By increasing the ISVP concentration to 300 pM, a greater number of ISVPs adhered to GUV membranes. Subsequently, these ISVP particles formed aggregates measuring a fewμm in size (Figure. 2 C, D) and induced more GUV rupture and membrane deformations. Interestingly, multiple GUVs, all exhibiting local membrane deformations, were bridged by the T1L ISVP aggregates (Figure 2 D-F). The junction between bridged GUVs was two membranes adhered together, not a single fused membrane, as their line-scan fluorescence intensity was twice the intensity of a single membrane (Figure. 2F). Moreover, the ISVP aggregates were confined at the intermembrane junctions (Figure. 2 E). In cases where multiple GUVs were bridged together into a cluster resembling foamy bubbles, the ISVP aggregates concentrated at the high-curvature junctions of the GUV membranes (Figure. 2 F). In occasional cases, the ISVP-induced membrane bridging eventually led to vesicle fusion (Figure. 2G; Video S4).

ISVP-to-ISVP* conversion results in the release of two types of peptides derived from the μ1 protein, μ1N and ϕ ^19-21^. These μ1-derived peptides are known to target membranes and form pores that are 4-9 nm in size ^13^, and μ1N alone is sufficient for pore formation ^20^. We therefore asked if the GUV membrane deformations were solely induced by these μ1-derived peptides ^13^. To test this, we prepared the μ1 peptides by converting 10 pM T1L ISVPs to ISVP*s via heat treatment. The particles were pelleted and the supernatant containing μ1 peptides was added to GUVs. Under these conditions, we found that even though the peptide-containing supernatant caused a small fraction (< 10%) of GUVs to rupture, it alone did not induce any membrane deformations similar to those that we observed in the presence of T1L ISVPs (Figure. S1), indicating that the membrane deformations require the presence of the ISVP capsids.

How did the ISVP capsid deform the lipid membranes? We reported previously that lipids facilitate the uncoating conversion of the metastable viral capsid from ISVP to ISVP* ^16^. Owing to the capsid structural changes, ISVP*s are expected to have a more hydrophobic surface than ISVPs and thus more likely to aggregate ^19^. Supporting this notion, we observed in control experiments that T1L ISVP*s generated *in vitro* from heat treatment formed large aggregates even in buffer solution, whereas the ISVPs remained monodisperse (Figure. S2). Therefore, the ISVP aggregates formed on the GUV membranes were likely to be partially uncoated ISVP*s. Their colocalization with lipids, as shown in Figure. 2, supports the idea that the semi-hydrophobic capsid of ISVP*s extracted lipids, via similar mechanisms by which large semi-hydrophobic nanoparticles extract lipids and disrupt lipid bilayers ^26^.

### Kinetics of viral capsid disassembly affect extent of membrane deformation

We next aimed to investigate whether the disassembly kinetics of the metastable ISVPs influence their interaction with the membranes. For this purpose, we employed the reovirus reassortant T1L/T3DM2. Structurally, T1L/T3DM2 differs from T1L in the M2 genome segment, which encodes the μ1 protein that converts to the membrane-active peptides μ1N and ϕ ^27^ (Figure. S3). The μ1 proteins of T1L and T1L/T3DM2 vary only in 9 of the 708 amino acids. Importantly, both virus strains have identical sequences of the pore-forming μ1N fragment. However, in comparison to T1L, T1L/T3DM2 ISVP uncoats more rapidly or to a greater extent ^27^. These differences enable T1L/T3DM2 to initiate infection more expeditiously in host cells ^28^.

If the membrane perturbations we have observed on GUVs are relevant to infection, we should expect that T1L/T3DM2 induces greater deformations in comparison to T1L. To test this idea, similar to the T1L ISVP experiments described above, we added 10 pM of fluorescently labeled T1L/T3DM2 ISVPs to the GUVs (DOPC, DOPE, 2:1 mol ratio, with 0.2% Bodipy-PE) at 37°C. We observed that T1L/T3DM2 ISVPs, like the T1L ISVPs, induced membrane deformations in most GUVs (Figure. 3A) and rupture of a small fraction of them. However, a few key differences indicated significantly stronger effect of T1L/T3DM2 ISVPs on lipid membranes. First, T1L/T3DM2 formed larger, more extensive aggregates than the T1L ISVPs, as they moved along the GUV membranes (Figure. 3A, B). Second, the T1L/T3DM2 ISVPs at 10 pM induced bridging between GUVs, which was only observed for 300 pM T1L ISVPs (Figure. 3B; Video S5). As we increased the concentration of T1L/T3DM2 ISVPs to 300 pM, they formed aggregates as large as tens ofμm and induced extensive membrane deformations (Figure. 3C, D). More rupture and bridging events of GUVs were captured during time-lapse epifluorescence imaging (Figure. 3 H, E, and F; Video S6, S7). Some GUVs ruptured after being bridged (Figure. 3 G; Video S8), suggesting a situation where the membrane perturbation could no longer be self-healed by the mobile lipid membrane. Thus, the greater disruption of membranes by T1L/T3DM2 matches the more efficient uncoating of its ISVP capsid during cell entry.

**Figure 3.**
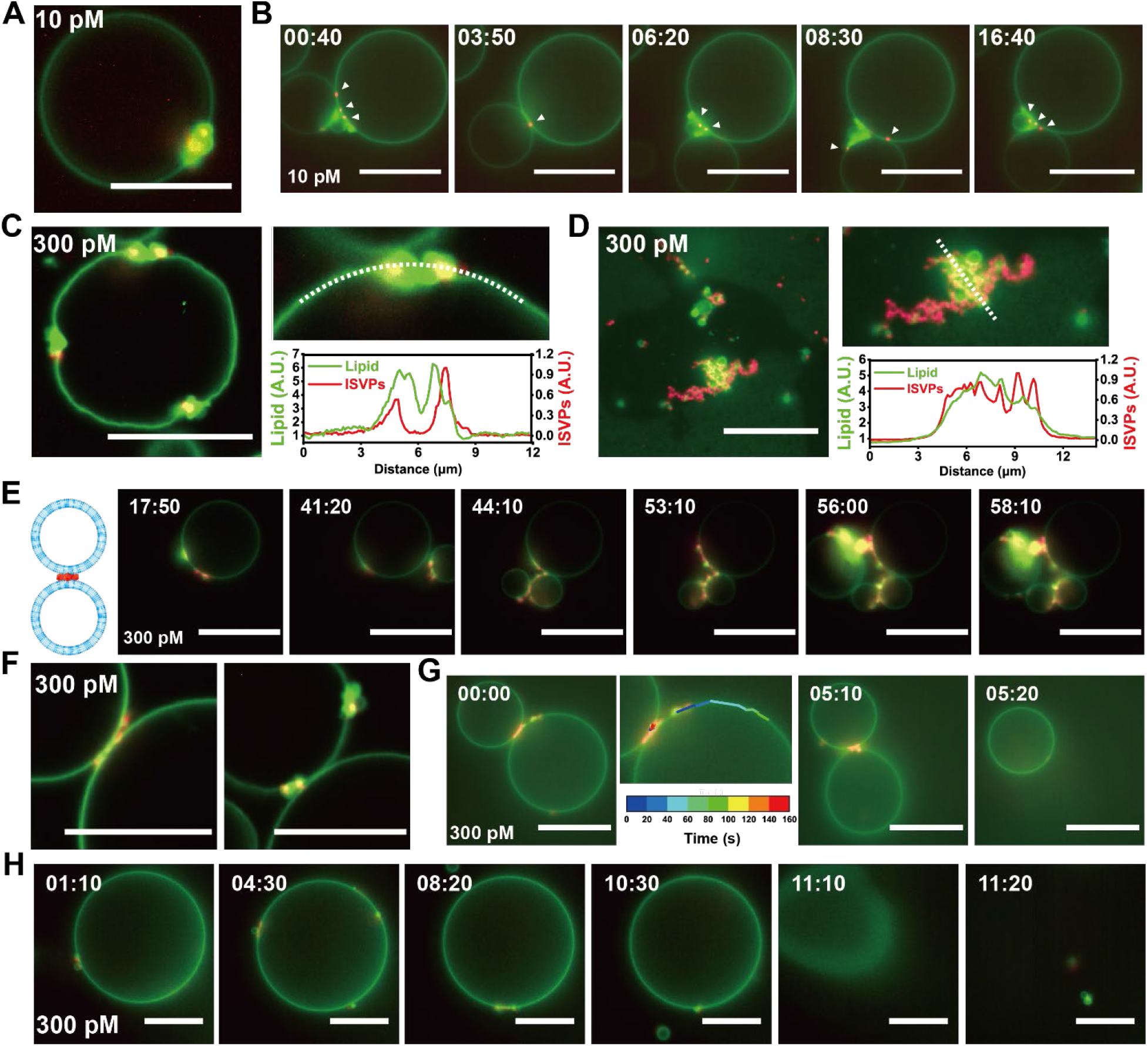
T1L/T3DM2 ISVPs induce membrane deformation, bridging, and rupture of GUVs. (A, B) Rescan confocal microscopy image (A) and epifluorescence time-lapse images (B) showing the adsorption and aggregation of T1L/T3DM2 ISVPs (shown in red) and the membrane deformation of a GUV (shown in green). ISVP concentration was 10 pM. (C) Rescan confocal microscopy images and line-scan plots showing the colocalization between T1L/T3DM2 ISVPs (300 pM) and membrane budding. (D) Epifluorescence images showing the colocalization of T1L/T3DM2 ISVP aggregates and lipids on a planar membrane formed from GUV rupture. ISVP concentration was 300 pM. (E) Schematic illustration and epifluorescence time-lapse images showing membrane bridging induced by 300 pM T1L/T3DM2 ISVP. (F) Rescan confocal microscopy images showing the bridging of GUVs by T1L/T3DM2 ISVPs (300 pM). (G) Epifluorescence time-lapse images showing GUV rupture after bridging induced by T1L/T3DM2 ISVPs (300 pM). The trajectories, color-coded with time, show the diffusion of some free ISVPs and confinement of some other ISVPs at the junction between bridged GUVs. (H) Epifluorescence time-lapse images showing GUV rupture induced by 300 pM T1L/T3DM2 ISVPs. The ISVPs colocalized with membrane budding and moved along the GUV surface. Scale bars in all images, 15 μm.

### Cholesterol enhances ISVP-induced membrane disruption

While model membranes comprised of PC and PE appeared sufficient to facilitate ISVP-to-ISVP* conversion, cholesterol levels are known to play a significant role in viral infections ^29^. For many viruses, such as the non-enveloped poliovirus and adenovirus, depleting cholesterol blocks cell entry ^30, 31^. It remains unclear how cholesterol affects reovirus interaction with membranes, but studies suggested that membrane cholesterol level influences either ISVP-to-ISVP* conversion or membrane disruption by reovirus ^32^. In our experiments, we prepared GUVs with varying amounts of cholesterol, from 0% (DOPC: DOPE = 2:1 molar ratio), 7.7 mol% (DOPC:DOPE: cholesterol = 8:4:1), 14.3 mol% (DOPC:DOPE: cholesterol = 4:2:1), to 25.0 mol% (DOPC:DOPE: cholesterol = 2:1:1). After adding 10 pM of either T1L or T1L/T3DM2 ISVPs to the GUVs, we observed more ISVPs adhered to the membranes containing cholesterol and formed larger aggregates compared to that on membranes without cholesterol. Subsequently, the ISVPs induced more GUV rupture, membrane deformations, and bridging between GUVs (Figure. 4 A). This suggests that cholesterol enhances the ISVP-membrane interactions, and the effect was most prominent in the presence of 7.7% and 14.3% cholesterol.

**Figure 4.**
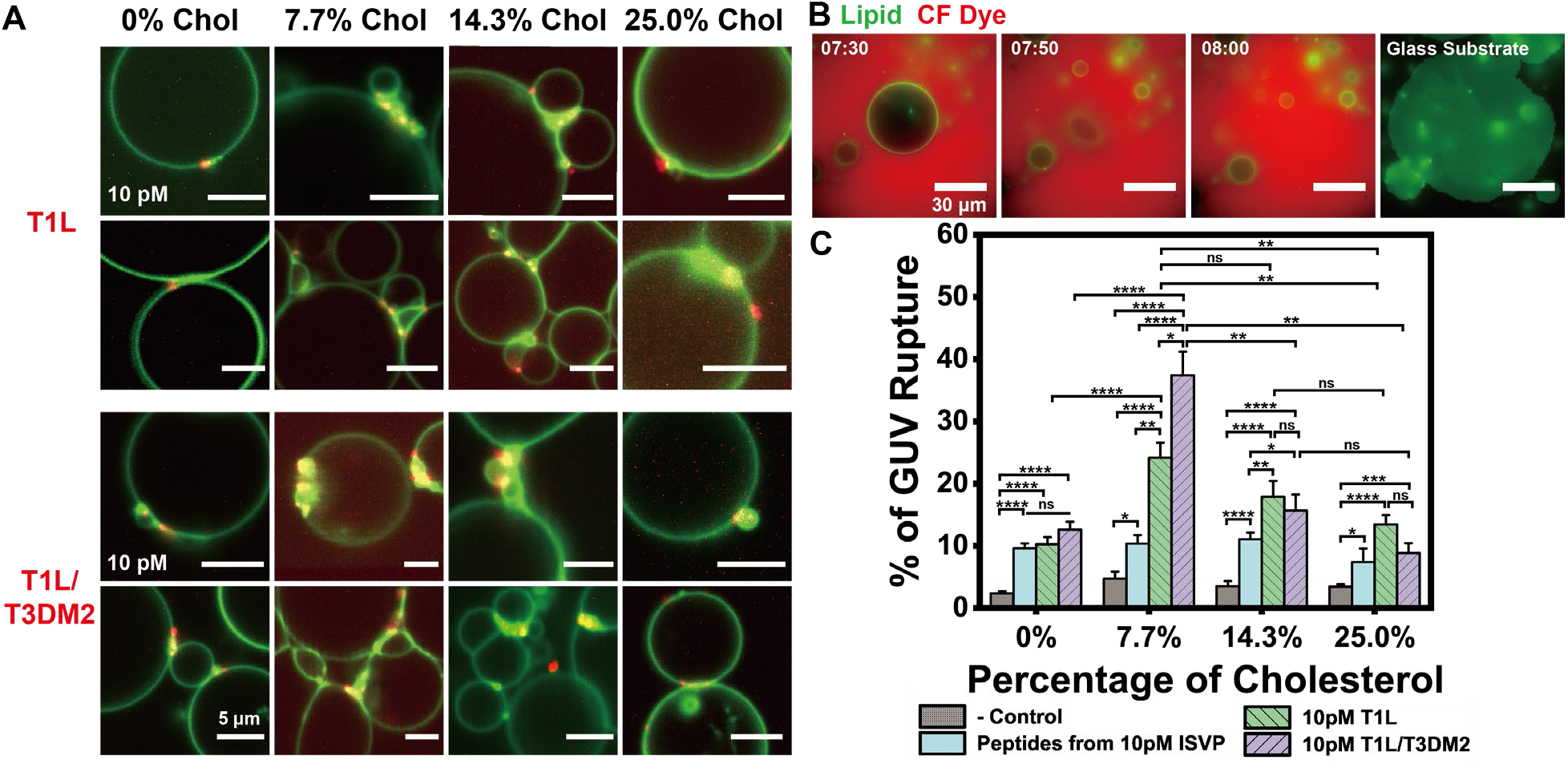
Cholesterol enhances ISVP-induced membrane deformation and rupture of GUVs. (A) Rescan confocal microscopy images showing the membrane deformation and bridging of GUVs of varying cholesterol content, induced by T1L ISVPs (10 pM) and T1L/T3DM2 ISVPs (10 pM) (shown in red). Scale bars in all images in (A), 5 μm. (B) Epifluorescence images showing the rupture of a GUV after ISVP interaction to form a planar membrane (shown in green). Scale bars in all images in (B), 30 μm. (C) Bar graphs showing the percentage of GUV rupture after adding 10 pM ISVPs or supernatant containing μ1-peptides. Statistical significance is highlighted by P values (Student’s t-test) as follows: **** P < 0.0001; *** P < 0.001; ** P < 0.01; * P < 0.05; ns P > 0.05, not significant. Error bars represent the standard error of the mean (S.E.M.).

To quantify the effect of cholesterol on GUV rupture, we developed a GUV rupture assay ^33, 34^. In this assay, after incubating ISVPs with GUVs for 30 min at 37°C, we imaged the planar membranes formed by ruptured GUVs on the coverslip bottom of imaging chambers and measured the membrane area (Figure. 4B, S4 A). We then measured the total membrane area of all GUVs in a control sample by intentionally rupturing all GUVs with a buffer solution of higher osmolarity (Figure. S4 C). We optimized the GUV concentration to ensure that only single sheets of bilayer were formed from the GUV rupture (Figure. S4 B, C, and D) and kept the GUV concentration the same for all samples. Under those conditions, the ratio of the membrane area from ISVP-ruptured GUVs to that of the total GUVs is equivalent to the fraction of GUVs ruptured by ISVPs. Consistent with our observation from confocal fluorescence imaging (Figure. 4A), cholesterol enhanced GUV ruptures induced by either T1L or T1L/T3DM2 ISVPs, compared to GUVs without cholesterol. Importantly, GUVs containing 7.7 mol% of cholesterol had the most significant rupture (Figure. 4C). This non-monotonic dependence of GUV rupture on the cholesterol content is qualitatively consistent with previous reports that either depletion ^30, 31^ or overly accumulation ^32^ of cholesterol blocked the infectivity of non-enveloped viruses. Further, we found that when the ISVP concentration was increased to 300 pM, both T1L and T1L/T3DM2 strains caused the rupture of nearly all GUVs containing 7.7 mol% cholesterol.

In contrast to the significant enhancement of ISVP-induced membrane rupture by cholesterol, its presence had no effect on GUV rupture induced solely by the supernatant containing μ1-derived peptides (Figure. 4C). ISVPs induced significantly more GUV rupture thanμ1 derived peptides in membranes with 7.7 mol% and 14.3 mol% cholesterol. This observation suggests that the majority of membrane rupture resulted from interactions between the ISVP capsid and lipids, rather than the pore-forming effect of releasedμ1 peptides. This comparison becomes more pronounced when considering that the peptide concentration in the supernatant generated from 10 pM ISVPs under heat treatment should be higher than that released from 10 pM ISVPs during the ISVP-membrane interaction, as heat treatment maximizes the disassembly of the ISVP capsid (Figure. S2 B). With this in mind, at 0% cholesterol, when ISVPs induced a similar fraction of GUV rupture as the μ1 peptides, some of the rupture events are expected to be caused by the ISVP capsids.

### Cholesterol enhances ISVP adsorption on membranes and formation of large pores

To further quantify the effect of cholesterol on ISVP-membrane interactions, we next measured and compared the kinetics of ISVP adsorption on membranes with and without cholesterol. To measure this on a curved GUV membrane was challenging. Instead, we prepared planar lipid membranes composed of either lipid composition: 2:1 DOPC: DOPE (0% cholesterol) or 8:4:1 DOPC: DOPE: cholesterol (7.7 mol% cholesterol) (Figure. 5 A). By imaging the adsorption of single T1L/T3DM2 ISVPs as a function of time, we observed that the inclusion of 7.7 mol% cholesterol in the planar lipid membrane increased the adsorption of ISVPs by about five times (Figure. 5 B). Interestingly, the number of ISVPs adsorbed on the membrane increased exponentially with time. This suggests a positive feedback mechanism where previously adsorbed ISVPs created new adsorption sites, possibly by releasingμ1 peptides that serve as membrane anchors for recruiting more ISVPs ^16, 21^.

**Figure 5.**
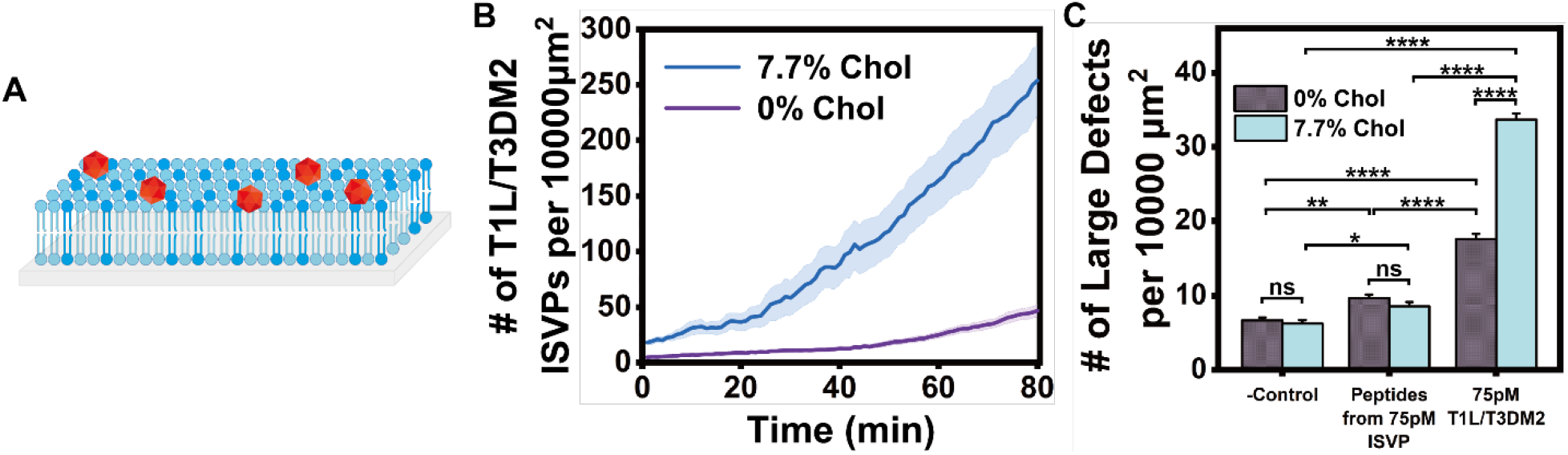
Cholesterol enhances ISVP adsorption to membranes and the formation of large pores. (A) Schematic illustration of ISVPs interaction with a planar-supported lipid bilayer. (B) Averaged line plots showing the surface density of T1L/T3DM2 ISVPs (10 pM) adsorbed on a planar lipid membrane containing different molar percentages of cholesterol as a function of time. Time zero was immediately after ISVPs were added to the membranes. Each line plot represents average ± S.E.M. from more than 6 samples. (C) Bar graphs showing the surface density of large pores formed on the planar lipid membrane containing different molar percentages of cholesterol, after a 30 min incubation with 75pM T1L/T3DM2 ISVPs or μ1 peptide-containing supernatant prepared from 75 pM ISVPs. Statistical significance is highlighted by P values (Student’s t-test) as follows: **** P < 0.0001; ** P < 0.01; * P < 0.05; ns P > 0.05, not significant. Error bar represents S.E.M.

Using the planar membranes, we further quantified the effect of cholesterol in pore formation by ISVPs. After T1L/T3DM2 ISVPs interacted with planar lipid membranes at 37 °C for 30 min, we washed away free ISVPs and then added fluorescently labeled bovine serum albumin (BSA) to the membrane. As BSA is known to adsorb strongly on bare glass surfaces but not on zwitterionic lipid bilayers, it backfilled pores in membranes where the glass surface was exposed ^35, 36^. The BSA-filled pores were then visualized using total internal reflection fluorescence (TIRF) microscopy and super-resolution radial fluctuations (SRRF) image postprocessing (Figure. S6) ^37^. Given that the dimensions of a BSA molecule are approximately 4 nm x 4 nm x 14 nm ^38^, this backfilling method can only detect membrane pores larger than the size of a BSA. Indeed, we did not detect the formation of large pores in membranes after adding ISVP* supernatant containingμ 1 peptides (Figure. 5 C), which are known to induce pores of 4-9 nm in size ^13^. By contrast, T1L/T3DM2 ISVPs induced large pores in 0% cholesterol membranes and significantly more in 7.7% cholesterol membranes. By measuring and comparing the fluorescence intensity of backfilled BSAs in individual pores to that of single BSAs, we estimated that an average of (20 ± 29) BSA molecules were inside each pore, which corresponds to an average pore diameter of (30 ± 19) nm (Figure. S6 E), calculated on the assumption of a globular shaped BSA (Figure. S6 B-D). Some pores were as large as 100 nm. These results indicated that while μ1-derived peptides can only generate small holes that are about 1/10 of the size of the reovirus particles, the metastable ISVP capsid facilitates and is required for pore expansion. Such an expanded pore can allow viral capsid to cross the membrane barrier and deliver the core into the cytoplasm for replication.

How does cholesterol affect the ISVP-membrane interactions? Previous studies have shown that cholesterol has a lipid-condensing effect, rendering unsaturated lipid bilayers less fluidic, stiffer, and thicker ^39, 40^. This lipid-condensing effect can be counterbalanced by adding the lipid 25-hydroxycholesterol (25-HC) in membranes ^41^. 25-HC is an oxysterol synthesized from cholesterol by the addition of a hydroxyl group at position 25-carbon of the isooctyl tail. Therefore, we next replaced cholesterol with 25-HC in GUV membranes to examine whether the lipid-condensing effect of cholesterol plays any role in the ISVP-induced GUV rupture. In experiments, we prepared GUVs containing 7.7 mol% 25-HC (DOPC: DOPE: 25-HC = 8:4:1 molar ratio) and added 10 pM T1L or T1L/T3DM2 ISVPs. Fluorescence confocal microscopy images and GUV rupture assay data showed that 25-HC inhibited ISVP adsorption on the membrane, resulting in negligible membrane deformations (Figure. S5 A). The fraction of ruptured GUVs was also substantially reduced to a level even lower than that induced byμ1 peptides alone (Figure. S5 B). The opposite effects of cholesterol and 25-HC on ISVP-induced GUV ruptures suggest that the lipid-condensing effect of cholesterol influences how ISVPs interact with and disrupt lipid membranes.

## Discussion

Non-enveloped viruses are known to release membrane lytic peptides during the entry process. The prevailing notion is that these peptides disrupt host cell membranes, facilitating viral entry. However, a paradox arises for many non-enveloped viruses, such as reovirus, where the pores induced by peptides in host membranes are too small for the viral capsid to traverse. Our previous studies, utilizing reovirus as a model, demonstrated that the presence of lipids promotes the structural disassembly of metastable intermediate infectious subviral particles (ISVP) into ISVP*s ^15, 16^.

In this study, we present direct microscopy evidence illustrating the dynamic process by which metastable ISVP capsids of reoviruses interact with lipids, deforming and permeabilizing host membranes to facilitate entry in both live cell and model lipid membrane systems. The distinctive steps in ISVP interaction with membranes leading to membrane disruption are revealed.

Initially, after adsorption to membranes, ISVPs do not remain stationary but diffuse along the membrane while remaining associated. As metastable ISVPs convert to ISVP*s, amphiphilic μ1 peptides are exposed on the capsid, and some are released. Based on previous work indicating that μ1 peptides, when inserted into the membrane, bind to and recruit the ISVP capsids to the membrane ^16^, the “surfing” motion of ISVPs is likely due to them being anchored to the membrane via exposed μ1 peptides on the viral capsid or those already released and embedded in the membrane. As ISVPs move along membranes, they gradually disassemble into ISVP*s with a more hydrophobic capsid surface, a feature that we confirmed from the aggregation of ISVP*s in aqueous buffers. Hydrophobic interaction becomes crucial in driving two concurrent steps following the ISVP “surfing” motion: the aggregation of uncoated ISVP capsids and the deformation of the membrane.

The ISVP-induced deformations include membrane budding, bridging, and, in some cases, rupture. These phenomena are strikingly similar to membrane deformations induced by semi-hydrophobic nanoparticles ^26^, supporting the central role of hydrophobicity in the process. Importantly, the extent of membrane disruption correlates positively with the infectivity of viral variants: capsids that disassemble more rapidly induce more pronounced membrane deformation and rupture.

Our results challenge prior notions by demonstrating that ISVPs can disrupt membranes upon conversion to ISVP*s. Previous studies had mainly focused on the role ofμ1 peptides. Earlier investigations have shown that ISVP-to-ISVP conversion induces membrane leakage in red blood cells^13, 14, 20^ and liposomes ^15, 16^. Pores thought to be caused byμ1 peptides were considered responsible for membrane penetration. In more recent studies, the release of macromolecules of finite sizes was utilized to provide more quantitative, albeit indirect, information about pore sizes. These studies suggested that the reovirusμ1 peptides induce pores ranging from 4 to 9 nm in size ^13^, which is significantly smaller than the size of an ISVP* or viral core. This left the puzzle unresolved: how do the viruses traverse the host membrane through pores that are too small? Our microscopy-based measurements directly visualize large membrane pores, indicating that the intermediate viral capsid particle formed during ISVP conversion to ISVP*, can generate pores significantly larger than those formed byμ1 peptides alone. We showed that the interaction between the capsid and lipid membrane is required for pore expansion and virus escape. As in endosomes, the release ofμ1-derived peptides is coupled with capsid-lipid interaction, the importance of the presence of capsid for perforation was overlooked previously. Furthermore, our results indicate that membranes of some vesicles are ruptured, while others are deformed. Initial membrane deformation can result in eventual rupture at a sufficiently high concentration of ISVPs. These data suggest that membrane deformation may occur prior to rupture. The inherent heterogeneity among virion particles may contribute to the subset of ISVP capsids capable of complete membrane rupture. It is known that not every reovirus particle is infectious but what leads to failure of some fraction of the particle is not known. Our data may suggest that, in some cases, it may be related to the inability to successfully penetrate the membrane.

Interestingly, during cell infection, we observed that viral particles move along the inner membrane of LAMP1-positive endolysosomes, and induce membrane bridging between adjacent endosomes. Because the ISVPs are in the endosomal lumen, it is unlikely that they directly serve as linkers to bridge multiple endosomes. Instead, we postulate that the bridging between endolysosomes is a result of membrane disruptions by the virus, based on previous reports indicating that lipid vesicles aggregate to stabilize membranes ^25^ with increased tension due to perturbation from peptides or nanoparticles ^42^. How such endosome aggregation contributes to viral infection is unclear. It is possible that fusion resulting from the endosome aggregations increases virus concentration in a single vesicular organelle, enhancing the chance of endosome escape. Whether the bridging interaction between endolysosomes is relevant to infection or aids in the repair of membranes breached by reoviruses remains to be determined in future studies.

Our study also reveals the influence of cholesterol on ISVP-membrane interactions. We demonstrate that cholesterol enhances ISVP adsorption onto membranes and the extent of membrane disruption, particularly at an intermediate cholesterol level. It is known that cholesterol levels play a significant role in viral infections ^29^. Depleting cholesterol was shown to block cell entry of non-enveloped viruses such as adenovirus and poliovirus ^30, 31^, while cholesterol accumulation in endosomes also impairs reovirus escape ^32^. It was proposed that membrane cholesterol level might influence either ISVP-to-ISVP* conversion or membrane disruption by reovirus ^32^. Our results provide a mechanistic explanation for those prior observations, although the exact cholesterol content used in the model membranes is not directly comparable to the range of cholesterol in endosome membranes. Further, we demonstrate that cholesterol influences ISVP-membrane interaction through its lipid-condensing effect. This conclusion is based on data illustrating that such an effect can be reversed by the addition of 25-HC, the oxidized analog of cholesterol. Previous work has indicated that 25-HC blocks fusion between enveloped viruses and host membranes, preventing the delivery of their genomic material into the cytoplasm ^43^. In the case of reoviruses, it has been demonstrated that the oxidation of cholesterol to 25-HC and accumulation of 25-HC in cells restricts the efficiency of cellular entry of reovirus virions ^44^. In line with these reports, our results collectively underscore the importance of cholesterol homeostasis in the host entry of non-enveloped viruses.

In summary, the model for reovirus-induced membrane penetration has proposed thatμ1-derived peptides released from metastable viral capsids facilitate membrane penetration ^13^. While true, our results elucidate the evolving metastable capsid’s role in this process. The observed dynamic interactions, membrane deformations, and disruptions contribute to a deeper understanding of how viral capsid-lipid interactions promote the entry of non-enveloped viruses. Additionally, the cholesterol dependence highlights the significance of cholesterol homeostasis in influencing the dynamics of virus infection. These insights provide valuable information for developing antiviral strategies and contribute to a broader understanding of virus infections in biological systems.

## Materials and Methods

### Materials and reagents

Joklik’s minimal essential medium was purchased from Lonza (Walkersville, MD). Fetal bovine serum (FBS) was purchased from Life Technologies (Carlsbad, CA). L-glutamine, Penicillin, and streptomycin were purchased from Invitrogen (Carlsbad, CA). Vertrel-XF specialty fluid was purchased from Dupont (Wilmington, DE). N -p-tosyl-L-lysine chloromethyl ketone (TLCK)-treated chymotrypsin was purchased from (Worthington Biochemical, Lakewood, NJ). FuGENE^®^ HD transfection reagent was purchased from Promega (Madison, WI). CF^®^568 succinimidyl ester (CF568-NHS) was purchased from Biotium (Fremont, CA). Dulbecco’s modified eagle medium (DMEM) and Alexa Fluor 647 succinimidyl ester (AF647-NHS) were purchased from ThermoFisher Scientific (Waltham, MA). 4-(2-hydroxyethyl)-1-piperazineethanesulfonic acid (HEPES), 5(6)-carboxyfluorescein, albumin from bovine serum (BSA), amphotericin B, phenylmethylsulfonyl fluoride (PMSF), and Amicon Ultra filters (30K) were purchased from MilliporeSigma (St. Louis, MO). LAMP1-GFP plasmid was kindly provided by Prof. Sergio Grinstein (University of Toronto, Ontario, Canada). 1,2-dioleoyl-sn-glycero-3-phosphocholine (DOPC), 1,2-dioleoyl-sn-glycero-3-phosphoethanolamine (DOPE), cholesterol, 25-hydroxycholesterol, 1,2-dioleoyl-sn-glycero-3-phosphoethanolamine-N-(lissamine rhodamine B sulfonyl) (RhB-DOPE), 1,2-dioleoyl-sn-glycero-3-phosphoethanolamine-N-[(dipyrrometheneboron difluoride)butanoyl] (TopFluor^®^-PE (Bodipy-PE)), and the mini-extruder were purchased from Avanti Polar Lipids, Inc (Alabaster, AL). ITO-coated slides (70-100 ohms, 25 mm × 50 mm × 1.1 mm) were purchased from Delta Technologies, Ltd (Loveland, CO). Ultrapure water (18.2 MΩ.cm) was used in all experiments.

All buffers used in the experiments: Dialysis buffer contained 10 mM Tris-HCl, 15 mM MgCl_2_, and 150 mM NaCl (pH 7.4). HEPES buffer contained 2 mM HEPES and 2 mM NaCl (pH 7.4). Glucose solution contained 100 mM glucose and 0.33 mM HEPES (pH 7.4). 2× glucose solution contained 200 mM glucose and 0.66 mM HEPES (pH 7.4). Sodium bicarbonate buffer contained 50 mM NaHCO_3_ (pH 8.5).

### Cells and viruses

Spinner-adapted murine L929 (L) cells were grown at 37 °C in Joklik’s minimal essential medium supplemented with 5% FBS, 2 mM L-glutamine, 100 U/mL penicillin, 100 μg/mL streptomycin, and 25 ng/mL amphotericin B. Reovirus type 1 Lang (T1L) and T1L containing the type 3 Dearing M2 gene segment (T1L/T3DM2) were generated by plasmid-based reverse genetics ^27, 45, 46^.

Vero 76 cells were cultured in DMEM complete medium supplemented with 10% fetal bovine serum, 2 mM L-glutamine, 100 U/mL penicillin, and 100 mg/mL streptomycin at 37 °C and 5% CO_2_. Transfection was carried out according to the manufacturer’s instructions. In brief, 0.125 million Vero 76 cells were seeded on a precleaned 30 mm glass coverslip without antibiotics 24 h before transfection. After replacing the cell medium with complete DMEM medium without antibiotics, 2 μL 500 ng/mL plasmid was then mixed with 3 μL FuGENE^®^ HD transfection reagents in 95 μL DMEM medium without supplements for 15 min. The mixture was then gently added to the cells. Cells were incubated overnight before use ^47^.

### Virus purification

Viruses were propagated and purified as previously described ^27, 48^. Briefly, L cells infected with second or third passage reovirus stocks were lysed by sonication. Virus particles were extracted from lysates using Vertrel-XF specialty fluid ^49^. The extracted particles were layered onto 1.2 to 1.4 g/mL CsCl step gradients. The gradients were then centrifuged at 187,000 × g for 4 h at 4°C in a SW 41 Ti rotor (Beckman Coulter, Brea, CA). Bands corresponding to purified virus particles (∼1.36 g/mL) were isolated and dialyzed into the dialysis buffer ^50^. Following dialysis, the particle concentration was determined by measuring the optical density of the purified virus stocks at 260 nm (OD260; 1 unit at OD260 = 2.1 ×10^12^ particles/mL) ^50^.

### Conjugation of reovirus to fluorescent dyes

Purified reovirus was labeled using CF568-NHS dye. Briefly, virions (1 ×10^13^ particles/mL) were diluted into fresh Sodium bicarbonate buffer and incubated with 10 μM CF568-NHS for 90 min at room temperature in the dark. After 90 min, the reaction was quenched by the addition of dialysis buffer, and the labeled virions were layered onto 1.2- to 1.4 g/mL CsCl step gradients and repurified as described above.

### Generation of infectious subvirion particles (ISVPs)

T1L or T1L/T3DM2 virions (2×10^12^ particles/mL) were digested with 200 μg/mL TLCK-treated chymotrypsin in a total volume of 100 μL for 1 h at 37°C. After 1 h, the reaction mixtures were incubated on ice for 20 min and quenched by the addition of 1 mM PMSF. The generation of ISVPs was confirmed by SDS-PAGE and Coomassie staining ^15^.

### Generation of ISVP* supernatant

The supernatant of pre-converted ISVP*s was generated as previously described ^20, 21^. Briefly, T1L or T1L/T3DM2 ISVPs (2 ×10^12^ particles/mL) were incubated at 52°C for 5 min. The heat-inactivated virus was then centrifuged at 16,000 × g for 10 min at 4°C to pellet particles.

### Fluorescence microscopy

All epifluorescence images and total internal reflection fluorescence (TIRF) images were acquired using a Nikon Eclipse Ti-E inverted microscope equipped with a Nikon 100×/1.49 N.A. Oil-immersion TIRF objective or a Nikon 40×/0.95 N.A. air-immersion objective and a Hamamatsu ORCA-Fusion digital CMOS camera. Re-Scan confocal microscopy (RCM) images were acquired using the RCM module (Confocal.nl, Netherland) added on the Nikon Eclipse Ti microscope with a Hamamatsu ORCA-Fusion digital CMOS camera. Glass coverslips for imaging were cleaned by sonication with 70% ethanol aqueous solution and rinsed with Milli-Q^®^ water before assembled into imaging chamber.

### Single-particle localization tracking

To track the movements of viral particles on vesicular or planar membranes, the centroids of single viral particles in epifluorescence or TIRF fluorescence microscopy images were localized using a radial-symmetry-based localization algorithm as previously reported ^51, 52^.

### Giant unilamellar vesicle (GUV) experiments

#### (a) GUV electroformation

Lipids of various compositions were mixed in chloroform to prepare the stock lipid solutions ^26^. 10 μL of lipid stock solution (5.0 mg/mL) was spread onto an ITO-coated glass slide to make a lipid film. Lipids were dried under nitrogen gas for over 30 minutes to remove residual chloroform. The lipid-coated ITO slide was immediately assembled with another ITO slide and a silicone spacer (1.7 mm thick) in a sandwich fashion and served as the electroformation chamber. The chamber has an approximate volume of 0.75 mL. 100 mM aqueous sucrose solution was added to the dried lipid film and vesicles were electroformed for 2 h under a sinusoidal alternating current (AC) field (3.4 V_rms_, 5 Hz). GUVs were used within two hours after electroformation. Pipette tips for transporting GUVs were truncated to avoid generating sheer stress.

#### (b) Imaging of GUV ISVPs interactions

GUVs were suspended in 100 mM glucose solution to a final lipid concentration of 20 μg/mL (assuming no loss of lipids during electroformation) before being added to an imaging chamber. This sucrose-glucose mixture helps GUVs settle to the bottom of imaging chambers before imaging ^26^. ISVPs suspended in glucose solution were added into the GUVs to reach a final ISVP concentration of either 10 pM (6×10^9^ ISVP/mL) or 300 pM (1.8×10^11^ ISVP/mL). The sample was imaged at 37 °C with 100× objective. Time-lapse multi-channel epifluorescence images or RCM fluorescence images were acquired to record the fluorescence emission of Bodipy-PE (Ex: 502 nm, Em: 511 nm) and CF568 (Ex: 562 nm, Em: 583 nm).

#### (c) Imaging of GUV dye influx

GUVs were suspended in 100 mM glucose solution containing 25 μM carboxyfluorescein in an imaging chamber ^26^. The final lipid concentration was 20 μg/mL. ISVPs of a final concentration of 300 pM (1.8×10^11^ ISVPs/mL) or peptide-containing supernatant generated from an equivalent amount of ISVPs were added to the GUVs. The sample was imaged with 100× objective at 37 °C. Time-lapse multi-channel epifluorescence images were acquired to record the fluorescence emission of RhB-DOPE (Ex: 560 nm, Em: 580 nm) and carboxyfluorescein (Ex: 493 nm, Em: 517 nm).

#### (d) GUV rupture assay

GUVs were suspended in 100 mM glucose solution in imaging chambers. The final lipid concentration was 8.87 μg/mL. ISVPs of a final concentration of 10 pM (6×10^9^ ISVP/mL) or peptide-containing supernatant generated from 10 pM ISVPs were added to the GUVs and incubated at 37 °C. For negative control experiments, an equivalent amount of dialysis buffer was added to the GUVs and incubated at 37 °C. For positive control experiments, GUVs were suspended in imaging chambers with the 2× glucose solution and then ruptured by mixing with the dialysis buffer at a 1:1 (v:v) ratio at 37 °C. After 30 min of interaction, the ruptured lipid bilayers deposited on the coverslip were washed 15 times with the glucose solution to remove unruptured GUVs ^33^. At each step of washing, truncated pipette tips were used to avoid disrupting unruptured GUVs, and only half of the solution was removed to avoid drying the lipid. The samples were imaged with 40× objective at room temperature. Epifluorescence images were acquired at random locations to record the fluorescence emission of Bodipy-PE. The images were intensity-thresholded by ImageJ and converted to binary masks, which were subsequently used to calculate the percentage of lipid surface coverage ^34^. The percentage of GUV rupture was calculated by normalization of lipid surface coverage with the positive control of each composition.

### Planar-supported lipid bilayer experiments

#### Large unilamellar vesicles (LUVs) formation

LUVs of 100 nm in diameter were made using the extrusion method ^35, 52^. Lipids with the desired composition were mixed in chloroform and dried in a round-bottom flask under nitrogen flow for over 30 min. Dried lipid films were hydrated in HEPES buffer to a final lipid concentration of 1 mg/mL. The lipid solution was vortexed and underwent freeze-and-thaw cycles six times before being extruded through a 100 nm filter membrane using a mini-extruder. The LUVs were stored at 4 °C.

#### (b) Preparation of planar-supported lipid bilayers

Planar lipid bilayers were formed via vesicle fusion on glass. Glass coverslips were sonicated in 70% ethanol aqueous solution and then in deionized water for 15 min each, cleaned in piranha solution (3:1 H_2_SO_4_ to 30% H_2_O_2_, volume ratio) for 15 min, and then rinsed with deionized water. The etched glass coverslip was assembled into an imaging chamber. To prepare supported lipid bilayers, LUVs were diluted with HEPES buffer to a final lipid concentration of 60 μg/mL and immediately added to the imaging chamber with a pre-etched glass coverslip bottom ^35, 52^. 1.5 mM CaCl_2_ aqueous solution was added to enhance vesicle fusion on glass coverslips. After incubation at room temperature for 40 min, the formation of lipid bilayer was complete, and excessive LUVs were rinsed away with HEPES buffer thoroughly. The supported lipid bilayer was blocked with 25 μg/mL BSA for 15 min and rinsed thoroughly with dialysis buffer.

#### (c) Quantification of ISVP adsorption on supported lipid bilayers

CF568 labeled T1L/T3DM2 ISVPs were added to supported lipid bilayers at a final concentration of 10 pM (6×10^9^ ISVP/mL) at 37 °C. TIRF microscopy time-lapse images were acquired at random locations for 80 min to record the fluorescence emission of CF568. Samples were kept at 37 °C in a temperature-controlled enclosure during imaging.

#### (d) Quantification of large pore formation in supported lipid bilayers

The first step was to prepare AF647-labeled BSA conjugates (AF647-BSA) for backfilling the large pores. To do this, 1 mg/mL BSA was mixed with 120 μg/mL AF647-NHS in sodium bicarbonate buffer (pH 8.5) and incubated under gentle rotation at room temperature for 2 h ^47^. Free dyes were then removed by centrifugation using Amicon 30K filters at 14000 × g at 4 °C for 10 min/wash for 6 times. Protein concentration and degree of dye labeling were measured using a ThermoFisher NanoDrop Nd-1000 microvolume spectrophotometer. The second step was the BSA backfilling assay. ISVPs of a final concentration of 75 pM, or peptide-containing supernatant generated from given concentrations of ISVPs (75 pM in this study), were added to the supported lipid bilayer and incubated at 37 °C for 30 min. After incubation, unbound ISVPs were removed by rinsing with dialysis buffer 6 times. Subsequently, AF647-BSA was added to the supported lipid bilayer at a final concentration of 8 μg/mL ^35^. After 15 min incubation, excessive AF647-BSA was removed by rinsing with dialysis buffer 15 times. TIRF microscopy images were acquired at random locations to record the fluorescence emission of AF647 (Ex: 650 nm, Em: 665 nm), which indicated the large defects on the supported lipid bilayer. Over 100 images were acquired consecutively at each location for the super-resolution radial fluctuations (SRRF) microscopy reconstruction ^37^.

### Live-cell fluorescence imaging of reovirus endocytosis

Vero 76 cells transfected with green fluorescence protein lysosomal associated membrane protein 1 (LAMP1-GFP) were incubated in a serum-free medium for 2 h and washed with PBS. CF568 labeled T1L or T1L/T3DM2 reovirus virions dispersed in pure DMEM were added to cells at a virion-to-cell ratio of 100,000: 1 and imaged at 37 °C. Time-lapse multi-channel TIRF microscopy images were acquired to record the fluorescence emission of CF568 and GFP (Ex, 482 nm; Em, 525 nm).

### Statistical analysis

Statistical evaluations of all data in this study were performed using Student’s t-test for two groups and one-way analysis of variance for multiple groups. Statistical significance is indicated as follows: ****p < 0.0001, ***p < 0.001, **p < 0.01, *p <0.05, and not significant (ns) p > 0.05. Statistic figures were plotted using OriginLab Pro 2023.

## Supporting Information

Supporting Information is available from the Wiley Online Library or the author.

## Acknowledgments

This work was supported by the National Institutes of Health (NIH) under award R21AI171911. The content is solely the responsibility of the authors and does not necessarily represent the official views of the National Institutes of Health. M.J. was partially supported by the Robert and Marjorie Mann Fellowship from Department of Chemistry, Indiana University. We thank Drs. Glenn Walpole and Sergio Grinstein (University of Toronto, Canada) for providing the LAMP1-GFP plasmid. The surface charge characterization of reovirus particles was done at the Nanoscale Characterization Facility at Indiana University.

## Author Contributions

M.J., P.D., and Y.Y. designed the research and wrote the manuscript. M.J. performed the experiments and analyzed the data. P.D. prepared and labeled reovirus particles.

## Data Availability Statement

The data that support the findings of this study are available from the corresponding author upon reasonable request.

## For Table of contents only

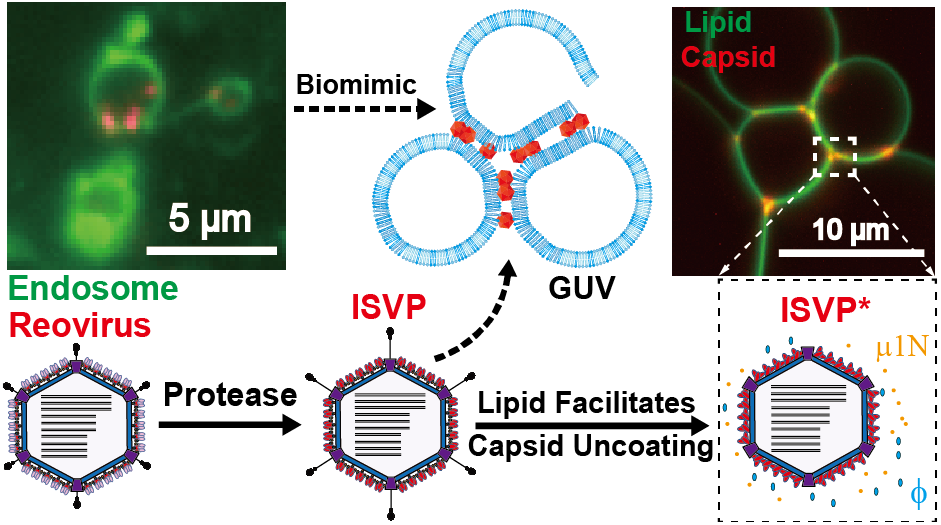

