## Supplemental Figures for "Cholesterol-Dependent Membrane Deformation by Metastable Viral Capsids Facilitates Entry"

**Contents**

Supplementary figures S1-S6

Supplementary videos S1-S3

A

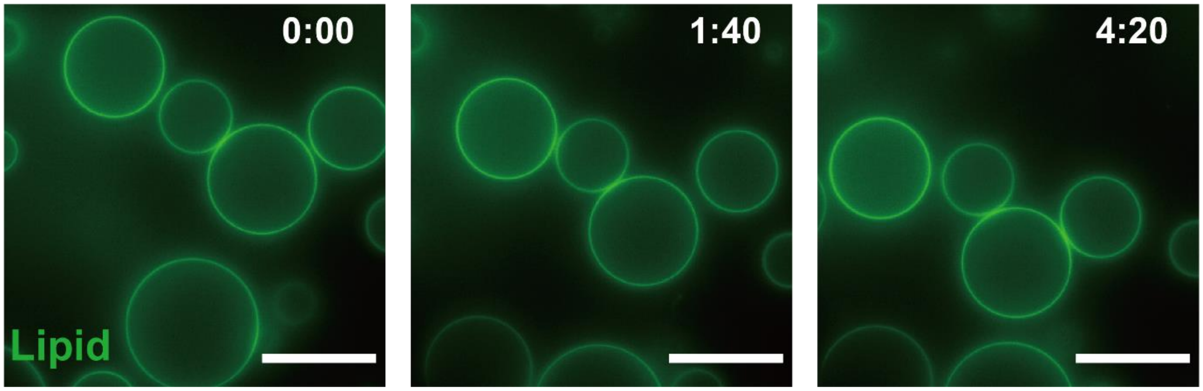


**Figure S1**. Effect of μ1 peptide on GUV membranes. Epifluorescence time-lapse images showing no membrane deformation of unruptured GUVs containing 0% cholesterol after the addition of μ1 peptides generated from 300 pM ISVPs. Scale bars in all images, 20 μm.

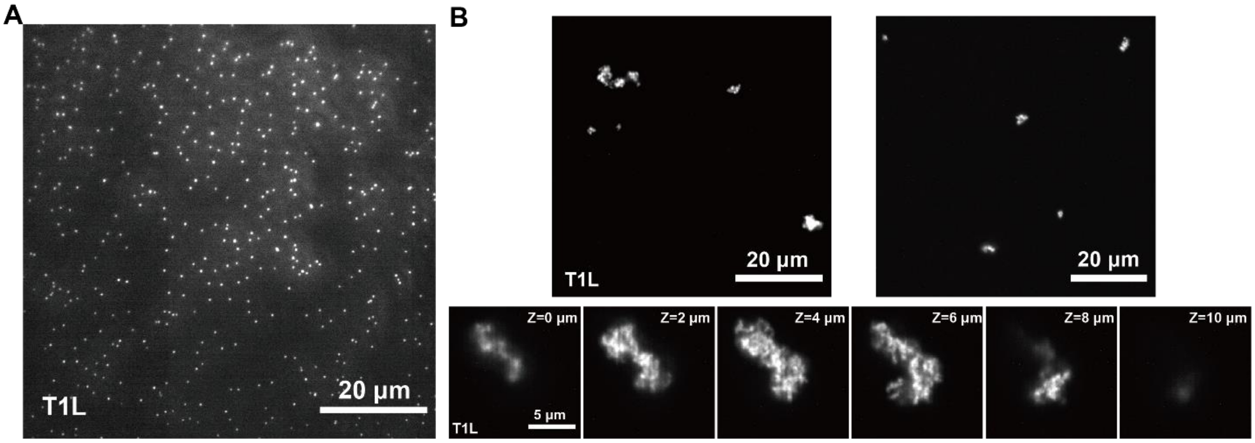


**Figure S2**. Monodisperse ISVPs and aggregations of ISVP*s in buffer. (A) TIRF microscopy images showing 10 pM CF568 labeled T1L ISVPs on the glass coverslip. (B) Rescan confocal fluorescence microscopy images showing 10 pM CF568 labeled T1L ISVP*s generated from heat treatment. (Top row) Images showing the ISVP* aggregates at different x-y locations of the same sample. (Bottom row) Z-stack images of the ISVP* aggregates.

A

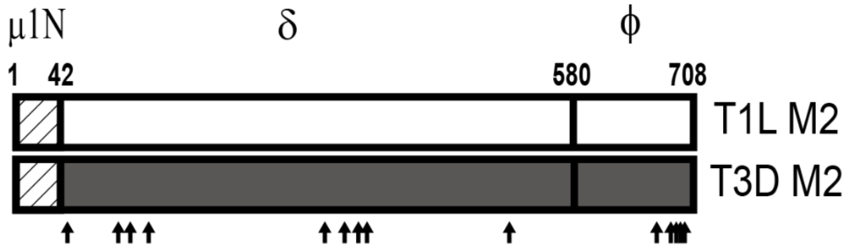


B

| Polymorphic Positions | Residue | |
|---|---|---|
| | T1L | T3D |
| 51 | I | V |
| 96 | N | I |
| 109 | A | T |
| 128 | L | M |
| 305 | V | A |
| 327 | K | R |
| 340 | T | S |
| 342 | G | N |
| 517 | E | D |
| 670 | T | A |
| 682 | N | S |
| 685 | G | S |
| 686 | S | G |
| 688 | P | S |
| 690 | A | N |

**Figure S3**. The difference in membrane penetration protein μ1 between T1L and T1L/T3DM2 strain. (A) Schematic diagrams of the μ1-encoding T1L and T3D M2 gene segments. The sequence of μ1N is identical for T1L and T1L/T3DM2 and is shown as a hatched box. The locations of polymorphic residues in T1L and T1L/T3DM2 μ1 are shown with arrows. (B) Location of polymorphic residues in T1L and T1L/T3D M2 μ1 proteins. The positions correspond to the arrows in (A).

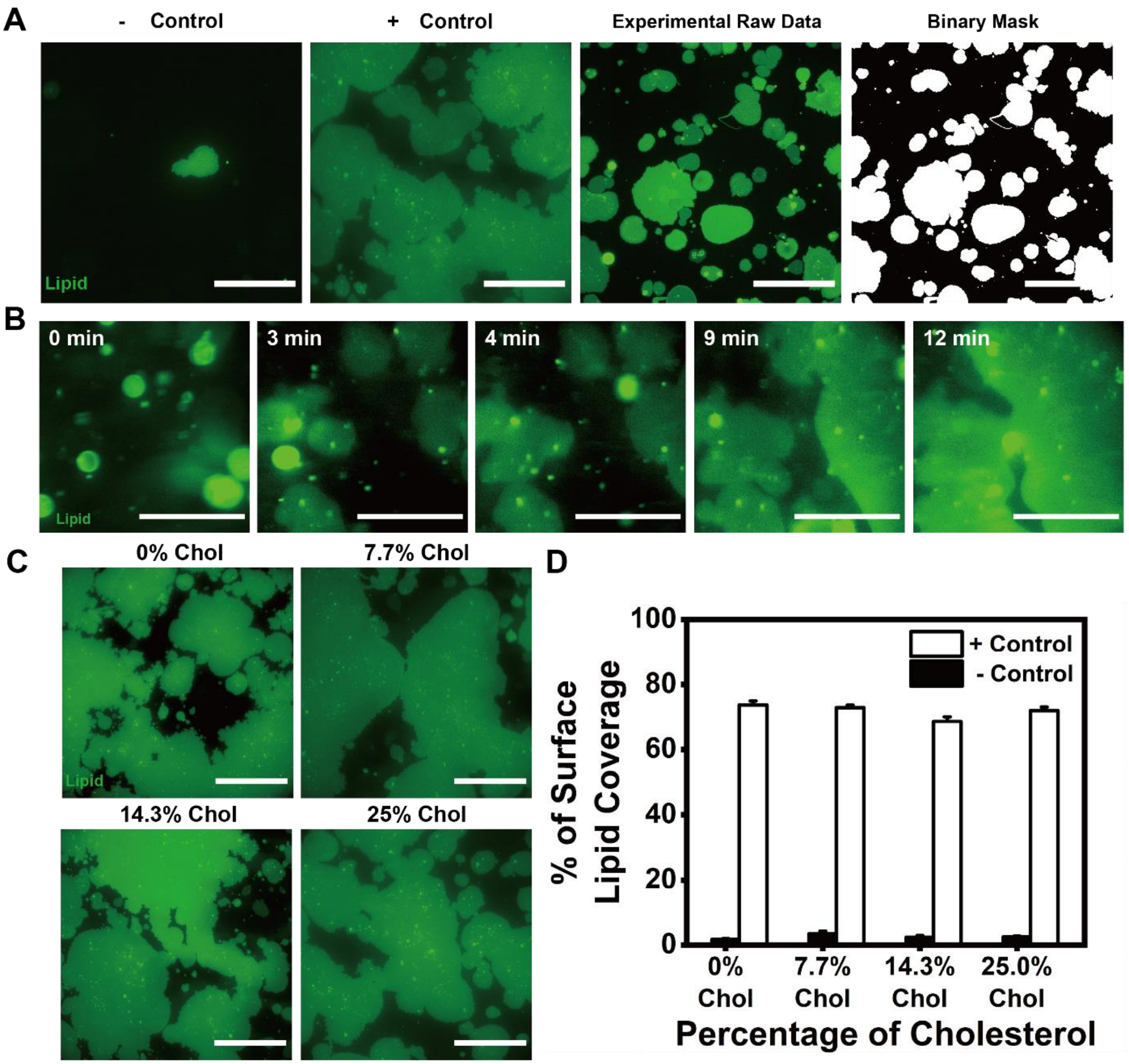


**Figure. S4** Quantification of the percentage of ruptured GUVs using a GUV rupture assay. (A) Epifluorescence images showing examples of the negative control (only GUVs), positive control (GUV + osmotic shock buffer), and experimental group (GUV + ISVPs or μ1 peptides). The raw images were converted to binary masks (white) and the percentage of surface areas occupied by the lipid membrane s was calculated using ImageJ. were calculated. The ratio of the lipid coverage percentage of experimental groups to that of positive controls was used to indicate the % of ruptured GUVs. (B) Examples of time-lapse epifluorescence images showing GUVs ruptured into planar lipid membranes. (C) Epifluorescence images showing examples of ruptured GUVs containing varying cholesterol content (mol%) in positive control experiments. (D) Bar graphs showing the percentage of lipid surface coverage from GUVs containing varying amounts of cholesterol in negative and positive control experiments. Error bars represent the standard error of the mean (S.E.M.). Scale bars in all images, 100 μm.

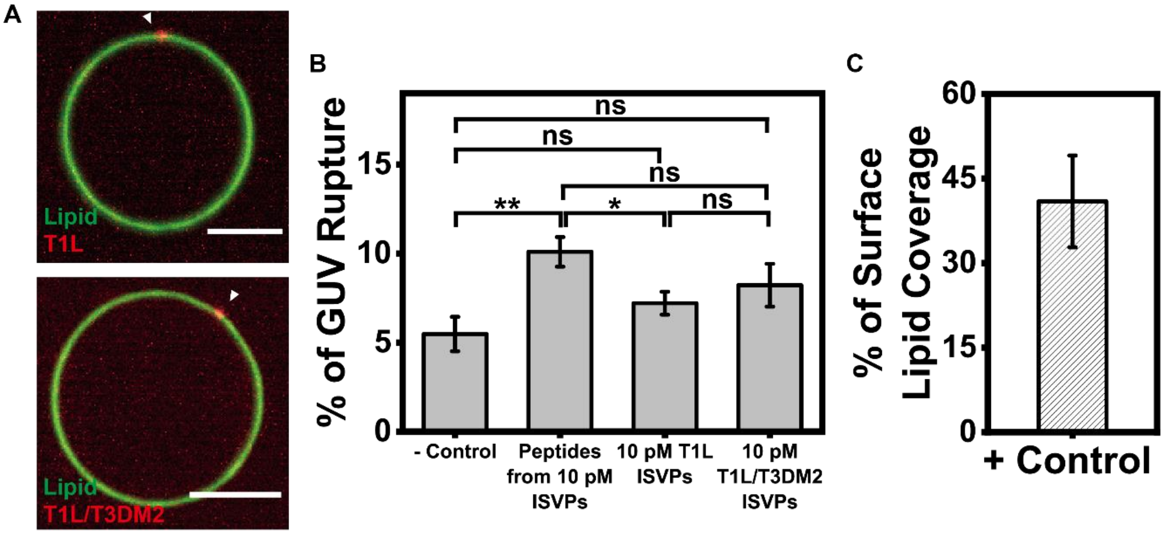


**Figure S5**. Effect of the addition of 25-hydroxycholesterol (25-HC) in GUVs on ISVP-membrane interaction. (A) Rescan confocal microscopy images showing no lipid disruption on GUVs containing 7.7 mol% 25-HC (DOPC, DOPE, 25-HC, 8:4:1 molar ratio) by T1L ISVPs and T1L/T3DM2 ISVPs at 10 pM ISVP concentration. Scale bars in all images, 5 μm. (B) Bar graphs showing the percentage of ruptured GUVs containing 7.7 mol% 25-HC under different experimental conditions as indicated. Statistical significance is highlighted by P values (Student's t-test) as follows: ** $P < 0.01$; * $P < 0.05$; ns $P > 0.05$, not significant. Error bars represent S.E.M. (C) Bar graph showing the percentage of lipid surface coverage of GUVs containing 7.7 mol% 25-HC in positive control experiments. Error bars represent S.E.M.

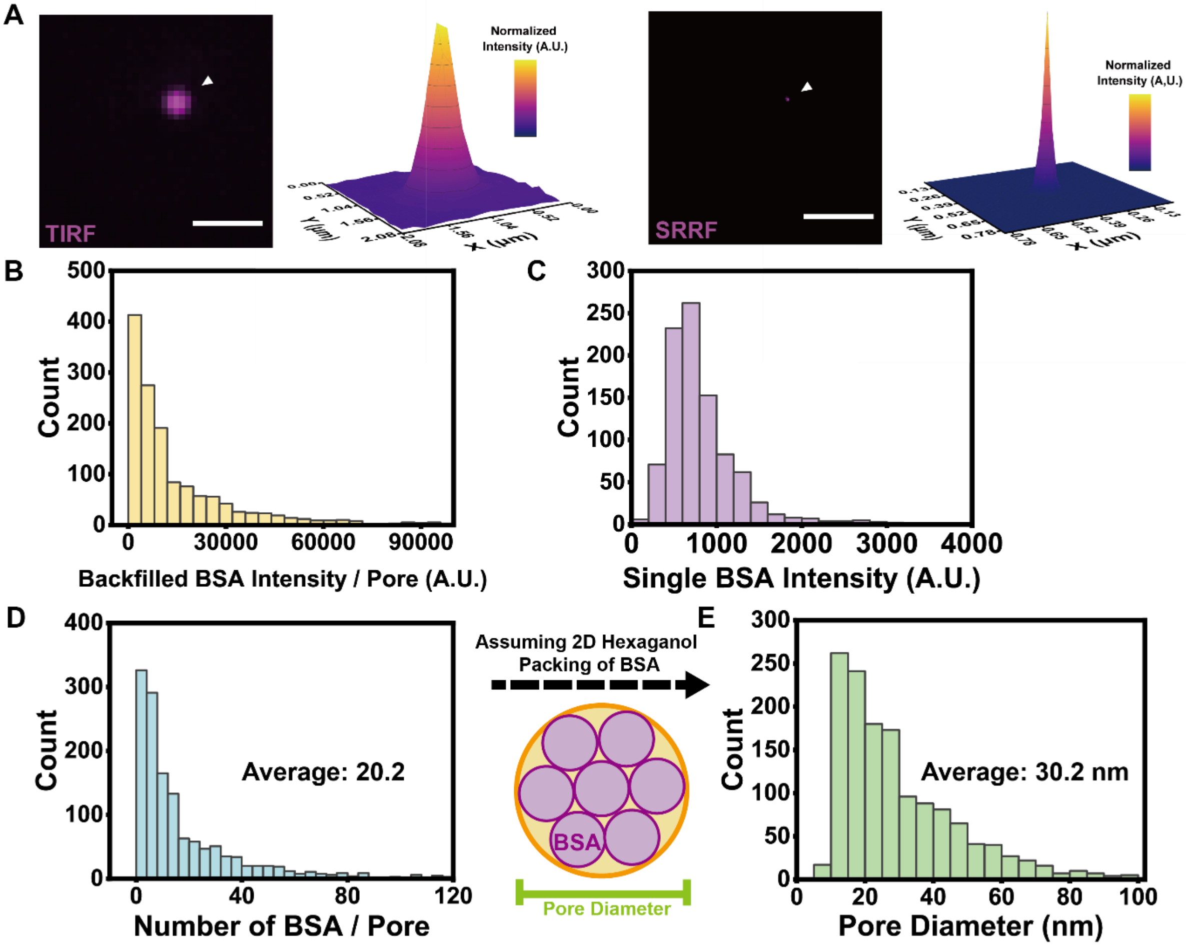


**Figure S6**. Quantification of large pores formed in planar lipid membranes. (A) TIRF microscopy image and super-resolution radial fluctuations (SRRF) microscopy image showing a large pore backfilled with AF647-BSA on the membrane. Scale bars in all images, 2 μm. (B) Histogram showing the fluorescence emission intensity of backfilled BSAs within large pores that were formed in planar lipid membranes containing 7.7 mol% cholesterol, after interaction with 75 pM T1L/T3DM2 ISVPs for 30 min (n = 1393 pores). (C) Histogram showing the fluorescence emission intensity of single BSA molecules on planar lipid membranes containing 7.7 mol% cholesterol (n = 943). (D) Histogram showing the estimated number of BSA molecules within each pore in planar lipid membranes containing 7.7 mol% cholesterol, after interactions with 75 pM T1L/T3DM2 ISVPs for 30 min (n=1393). (E) Histogram showing the estimated diameter of pores in planar lipid membranes containing 7.7 mol% cholesterol, after interactions with 75 pM T1L/T3DM2 ISVPs for 30 min (n=1393).

**Video. S1** Diffusion of T1L/T3DM2 ISVPs on the inner leaflet of an endosome in Vero cells.

TIRF fluorescence microscopy time-lapse video showing the diffusion of T1L/T3DM2 ISVPs (shown in red) on the inner leaflet of an endosome labeled with LAMP1-GFP (shown in green) in a Vero cell.

**Video. S2** Aggregation of endosomes containing T1L/T3DM2 ISVPs in Vero cells.

TIRF fluorescence microscopy time-lapse video showing the transport and aggregation of endosomes containing T1L/T3DM2 ISVPs (shown in red). Endosomes were labeled with LAMP1-GFP (shown in green) in Vero cells.

**Video S3.** T1L ISVPs induce membrane deformation while moving along the surface of a GUV.

Epifluorescence time-lapse video showing the adsorption and diffusion of T1L ISVPs (10 pM) on a GUV, resulting in membrane deformation.

**Video. S4** T1L ISVPs induce membrane fusion between GUVs.

Epifluorescence time-lapse video showing the bridging, fusion, and rupture of GUVs (shown in green) induced by T1L ISVPs (300 pM, shown in red).

**Video. S5** T1L/T3DM2 ISVPs induced deformation and bridging of GUVs.

Epifluorescence time-lapse video showing the disruptions and bridging on GUVs (shown in green) induced by 10 pM T1L/T3DM2 ISVPs (shown in red).

**Video. S6** T1L/T3DM2 ISVPs induced complete rupture of GUVs.

Epifluorescence time-lapse video showing the rupture of 2 GUVs (shown in green) induced by 300 pM T1L/T3DM2 ISVPs (shown in red).

**Video. S7** T1L/T3DM2 ISVPs induced briding of GUVs.

Epifluorescence time-lapse video showing the real-time bridging of GUVs (shown in green) induced by 300 pM T1L/T3DM2 ISVPs (shown in red).

**Video. S8** GUV rupture after briding by T1L/T3DM2.

Epifluorescence time-lapse video showing the briding of GUVs (shown in green) and subsequent rupture induced by 300 pM T1L/T3DM2 ISVPs (shown in red).
